# Comparative Analysis of De Novo Assemblers and Quantification Software for RNA-sequencing Data in Non-Model Arthropods

**DOI:** 10.1101/2025.08.01.668104

**Authors:** Marie V. Brasseur, Florian Leese, Christoph Mayer

## Abstract

**Background:** RNA-sequencing has greatly improved our understanding of the transcriptomic regulation of fundamental biological processes. Although the method has matured significantly within the last decade, bioinformatic processing of the resulting high-dimensional data sets is still challenging and the performance of algorithms can vary between data sets. As a consequence, for most non-model organisms, in particular arthropods, there is no or limited literature evidence which software is best suited to handle taxon-specific data characteristics. Therefore, we evaluated the performance of different *de nonvo* transcriptome assembler (Trinity, rnaSPAdes, IDBA-tran) and transcript quantification software (RSEM, Salmon) on transcriptomic data of a non-model insect and freshwater crustacean species, as well as the impact of different quality trimming strategies on the downstream bioinformatic processing results.

**Results:** While the trimming strategy had no considerable effect on the quality of transcriptome assemblies, the choice of the assembler had a substantial impact. IDBA-tran was less sensitive than the two other assemblers and produced the most fragmented transcriptome assemblies. The low remapping rates of reads against IDBA-tran assemblies further suggest that the input read data was not effectively leveraged by this algorithm. In contrast, Trinity and rnaSPAdes both generated comprehensive and contiguous *de novo* transcriptome assemblies, although Trinity appeared to be slightly more sensitive. This increased sensitivity, however, was associated with a higher redundancy in Trinity-generated assemblies compared to assemblies produced with rnaSPAdes. When the quality of the transcriptome assembly was high, RSEM and Salmon were able to identify the origin of at least 90% of the read data in the reference. Despite their different underlying quantification approaches, the estimated transcript counts of both tools were highly correlated and their expression signal was consistent. Notably, the alignment-free quantification algorithm Salmon was substantially faster than the alignment-based approach of RSEM. Furthermore, it was also slightly more sensitive, increasing the average re-mapping rate to ∼98%.

**Conclusion:** Since the performance of bioinformatic algorithms, especially of *de novo* assemblers, varies for different RNA-sequencing data sets, establishing an appropriate analysis workflow remains an important task. Our results show that the better performing combinations of algorithms produce congruent count data sets with consistent expression signal, highlighting the robustness of RNA-sequencing data analysis software.

## Introduction

Massively parallel sequencing of complementary DNA (cDNA), termed RNA-sequencing or RNA-seq, has revolutionized gene expression research (Wang et al., 2009). Compared to hybridization-based analysis methods such as microarrays, RNA-seq can profile all RNA molecules present in a biological sample at single-base resolution without any prior knowledge about the underlying genome (Finotello & Di Camillo, 2015; Wang et al., 2009). This key advantage and the development of sophisticated bioinformatic algorithms has made transcriptomic profiling accessible for non-model organism research (Ekblom & Galindo, 2011; Geniza & Jaiswal, 2017).

Yet, RNA-seq produces high-dimensional data sets that are often not easy to analyze (Conesa et al., 2016; Eldem et al., 2017). The properties of data sets from different experiments and organisms can vary due to technical (e.g., different library preparation methods, sequencing platforms, differences in sequencing coverage) and biological reasons (e.g., organism-specific differences in GC content, transcriptome complexity, levels of heterozygosity and genome repetitiveness) (García-Nieto et al., 2022; Shi et al., 2021). Despite the heterogeneity of expression data sets, bioinformatic algorithms are usually only assessed based on model organism or simulated data sets, for which the expression signal is known, but which are less complex and noisy than real biological data (see e.g., Bushmanova et al., 2019; Chandramohan et al., 2013; Teng et al., 2016; Voshall & Moriyama, 2018). In fact, benchmarking experiments involving animal taxa typically rely on extremely well-characterized models, such as human, mouse and fruit fly, whereas the algorithmic capacity to handle RNA-seq data from less studied taxa, in particular the hyperdiverse group of arthropods, is rarely assessed. Due to the (genomic) diversity of arthropods, the output of the different computational steps must be evaluated in order to create customized analysis workflows, as no algorithm fits all data sets (Hölzer & Marz, 2019; Kanitz et al., 2015; Teng et al., 2016). Therefore, we here evaluated the performance of different bioinformatic tools used to generate count data sets for differential gene expression (DGE) analyses from RNA-seq reads of non-model arthropods. We expect our study to support other researchers studying non-model organisms in selecting and establishing a suitable bioinformatic workflow. Since especially beginners in the field of transcriptomics require assistance in selecting an appropriate data analysis strategy, we also briefly outline the computational challenges inherent to the analysis of short-read RNA-seq data, how these challenges are addressed by the evaluated algorithms, and how the user can assess the quality of the generated output.

### RNA-Seq Data Analysis For Gene Expression Studies

If the aim of a study is to infer DGE patterns in a non-model species, a standard bioinformatic processing pipeline for RNA-seq data comprises three main steps prior to statistical testing:

1. Pre-processing of raw sequencing reads, i.e., quality control.
2. Reconstruction of a transcriptomic reference, i.e., *de novo* transcriptome assembly.
3. Transcript abundance estimation.

Reconstructing a representative transcriptome (step 2) and accurately quantifying transcript abundances (step 3) are fundamental prerequisites for a robust DGE analysis. Both steps are computationally challenging as they rely on relatively short sequencing reads (∼ 75-150 bp), compared to the length of the original biological sequences they are derived from (Grabherr et al., 2011; Li et al., 2010), e.g., average mRNA sequence lengths of 3,058 bp in *Drosophila melanogaster* (Adams et al., 2000). Further, the quality of the input data (step 1) is fundamental to ensure accurate downstream analysis results (Fabbro et al., 2013). While the implementation of quality trimming is computationally straightforward, the best trimming strategy for RNA-seq data is up to debate (Fabbro et al., 2013; MacManes, 2014; Williams et al., 2016). Therefore, we pre-selected three transcriptome assemblers and two quantification tools and evaluated their performance on Illumina RNA-seq data from two non-model arthropod species, and assessed the impact of two different quality trimming strategies (instead of different trimming software) on the downstream analysis results. The programs were chosen due to their high performance reported in the literature and the diversity of the underlying algorithms. Below we describe further details on aspects that are important for steps 1-3.

#### Step 1: Quality Control

Quality control is important to account for technical artifacts in raw sequencing reads. The most common error of Illumina platforms are nucleotide substitutions, which typically accumulate towards the 3’ end of the read (Fox et al., 2014; Stoler & Nekrutenko, 2021). The systematic error distribution is exploited by widely-used trimming tools like Cutadapt (Martin, 2011) or its wrapper script TrimGalore! (https://github.com/FelixKrueger/TrimGalore), which start at the 3’ end of the read and apply a quality cut-off when an increase in base quality is detected (see Cutadapt manual).

Another frequently observed technical artifact in sequencing data are homopolymers at the end of the reads (Schröder et al., 2010). In RNA-seq data sets, these homopolymers arise due to, e.g., mRNA enrichment using oligo(dT)-primers, which result in mononucleotide stretches of adenine (A) or thymine (T) at the end of the read (Pickrell et al., 2010). Additionally, libraries sequenced on Illumina NovaSeq or NextSeq platforms are biased towards high confidence overcalling of the guanine (G) base due to their two-color chemistry: during base calling, the absence of a fluorescence emission signal is interpreted as incorporation of a G (Van Pelt-Verkuil et al., 2019). In case of a premature termination of the read extension during sequencing, high-quality but erroneous homopolymer stretches are appended to the read (Andrews, 2016). If these Gs at the 3’ end mask the quality profile leveraged by trimming tools, they might hamper effective trimming of lower quality bases (Andrews, 2016). Therefore, we tested if removing low-complexity regions in the reads prior to quality trimming improves the outcome of downstream processing steps such as the *de novo* transcriptome assembly. For this, a C++ program was developed to remove homopolymers at the end of sequencing reads (for further details, see Supplementary Material S1).

#### Step 2: De Novo Transcriptome Assembly

If no reference genome or transcriptome exists, a transcriptome must be assembled *de novo* from the cleaned read data (Conesa et al., 2016; Geniza & Jaiswal, 2017). While the computational challenge inherent to short read data is not specific to transcriptome assemblies, the following biases unique to expression data must be addressed specifically: the coverage of different transcripts varies by several orders of magnitude and even different isoforms resulting from alternative splicing can show highly dynamic expression patterns (Grabherr et al., 2011). Shared sequence information, e.g., between alternatively spliced isoforms, introduces ambiguity, which is difficult to resolve (Grabherr et al., 2011). Many assembly algorithms were developed (e.g., Bushmanova et al., 2019; Grabherr et al., 2011; Peng et al., 2013) and their performance was evaluated extensively (e.g., Clarke et al., 2013; Hölzer & Marz, 2019; Voshall & Moriyama, 2018; S. Wang & Gribskov, 2017). Although these comparative studies consistently concur that there is no single optimal algorithm for all data sets, the Trinity (Grabherr et al., 2011) and rnaSPAdes (Bushmanova et al., 2019) assemblers are typically among the best performing programs identified therein. However, both assemblers can demand substantial computational resources, i.e., time and memory requirements (Hölzer & Marz, 2019; Lu et al., 2013; Zhao et al., 2011). The assembler IDBA-tran (Peng et al., 2013) instead requires significantly less memory than the two other programs and was shown to be generally fast, but performed only mediocrely (Hölzer & Marz, 2019).

Since performance but also computational efficiency are important parameters to consider in the analysis of complex data sets, we included all three programs in this benchmark study. Further information about the key differences between the three different algorithms, which are likely to affect their performance, are given in the Supplementary Material S2.

#### Step 3: Quantification

Identifying the true origin of RNA-seq reads in a transcriptomic or genomic reference is a non-trivial task. Ideally, the perfect algorithm would resolve the true loci of which all reads were derived, but repetitive regions, sequencing errors and in particular shared sequence information between paralogs, alternatively spliced isoforms and antisense transcripts introduce ambiguity (Li et al., 2010). Consequently, a substantial number of short reads map equally well to multiple locations (i.e., multi-mapping reads) in the reference (Li et al., 2010; Mortazavi et al., 2008).

Inferring transcript abundances comprises two steps: identifying the source of all transcriptomic reads in a given sample, followed by transcript abundance estimation (Li & Dewey, 2011; Nicolae et al., 2011; Patro et al., 2017). Two computational strategies exist to target the first part of the quantification problem: alignment-based (e.g., Li & Dewey, 2011; Nariai et al., 2014; Nicolae et al., 2011) or alignment-free (e.g., Bray et al., 2016; Patro et al., 2017) algorithms. Alignment-based algorithms are typically very accurate, but their precise read alignments in a base-per-base fashion require substantial computing times when the data set is complex (Bray et al., 2016; Kanitz et al., 2015). Alignment-free algorithms are substantially faster (Bray et al., 2016; Patro et al., 2014; Zhang et al., 2017); they exploit the idea that an approximation of the read origin is sufficient, termed mapping, if the aim is DGE quantification (Bray et al., 2016; Zhang et al., 2016) and search for a partial but exact overlap between a read and the reference (Bray et al., 2016; Patro et al., 2017; Srivastava et al., 2016).

The program RSEM is guided by read alignments and is often the best performing algorithm in benchmarking studies (Bray et al., 2016; Kanitz et al., 2015; Teng et al., 2016; Zhang et al., 2017). Despite being very accurate, RSEM is slow (Kanitz et al., 2015; Zhang et al., 2017). In contrast, the mapper Salmon was shown to be substantially faster and almost as accurate as RSEM (Teng et al., 2016; Zhang et al., 2017). Therefore, we included RSEM as representative for an alignment-based quantification software and Salmon as representative for an alignment-free software in this evaluation. Further information about the differences between the two quantification tools can be found in Supplementary Material S3.

## Methods

Two RNA-seq data sets generated in our lab were used for the comparative analysis of *de novo* assemblers and quantification software. These data sets represent transcriptomic data from the crustacean *Gammarus fossarum* (Amphipoda) (Brasseur et al., 2022), and from the insect *Ephemera danica* (Ephemeroptera) (Brasseur et al., 2023). The libraries of both data sets contain pooled RNA extracts of two (*G. fossarum*) or three (*E. danica*) specimens and were generated with different library preparation protocols and sequencing strategies (Table 1). A detailed description of the library preparation protocols is given in the respective studies.

**Table 1:** Library preparation strategies of the RNA-seq data sets used in this benchmarking study.

|  | *G. fossarum* | *E. danica* |
| --- | --- | --- |
| No. of libraries | 32 | 36 |
| RNA extraction | TRIzol + QIAGEN columns | GITC + magnetic beads |
| Library preparation | mRNA enrichment; unstranded | mRNA enrichment; stranded |
| Sequencing platform | HiSeq X | Illumina NovaSeq 6000 |
| Read length | 150 bp (paired-end) | 150 bp (paired-end) |

### Bioinformatic Processing

Quality of raw sequencing read data was checked with FastQC (Andrews, 2010). Homopolymers were trimmed with our custom C++ program Polyprune. Adapter removal and trimming of low-quality bases was performed with the Cutadapt v3.2 wrapper script TrimGalore! v0.6.6 in paired-end mode, applying a base quality cutoff Phred value of 20 and retaining only reads with a minimum length of 25 bp. The length filtering ensured to supply the same read data set to all assemblers, because Trinity only considers reads with a minimum length of 25 bp due to a fixed k-mer 25 (for further information, see Supplementary Material S2). *De novo* transcriptomes were assembled with Trinity v2.9.0 and v2.13.3, rnaSPAdes v3.15.0 and IDBA-tran v1.1.3. All assemblers were run in paired-end mode. Assemblies of *E. danica* were generated in strand-specific mode, except using IDBA-tran, which has no specific mode for stranded data. For the other parameters, the default settings were used, except for parameters specifying memory consumption and multi-threading. In total, six assemblies were produced for each species. Per species, two assemblies were generated with the same assembler, using either RNA-seq reads that were homopolymer trimmed prior to quality trimming or RNA-seq reads which were only subjected to quality trimming.

The cleaned sequencing reads used to generate the individual assemblies were either mapped with Salmon v1.9.0 or aligned with bowtie2 v2.3.5.1 (Langmead & Salzberg, 2012) followed by quantification with RSEM v1.3.3. Salmon was run in default mode except for parameters specifying different library preparation protocols (unstranded *G. fossarum* data: --libType IU; stranded *E. danica* data: --libType ISR) and parameters correcting for sequence-specific mapping biases (--validateMappings, --seqBias, --gcBias) as well as parameters controlling multi-threading. Similarly, RSEM was run with default parameters except the ones specifying multi-threading, paired-end data (--paired-end) and in the case of *E. danica*, strandedness (--forward-prob 0).

In total, 12 species-specific data sets were generated to evaluate the impact of the trimming strategy and the choice of the assembly and quantification software.

### Metrics to Evaluate the Performance of the Assembly and Quantification Programs

The quality of the transcriptome assemblies was evaluated based on predefined metrics, covering quality measures based on reference-free and reference-dependent metrics. Reference-free metrics refer to technical aspects of the assembly, including basic statistics (min., max., mean, median contig length), remapping rates of the reads used to assemble the transcriptome and the ExN50 statistic. The ExN50 value, proposed by the Trinity developers, is a modified version of the N50 value that is limited to the topmost highly expressed transcripts accounting for x% of the total expression (for further details, see the Trinity GitHub repository). These metrics were determined with scripts provided with Trinity and bash shell commands.

While useful, reference-free metrics cannot inform about biologically correct sequence composition, as opposed to reference-dependent metrics (Eldem et al., 2017; Voshall & Moriyama, 2018). For non-model organisms without genomic resources, the gene content of an assembly can be assessed using homology searches (Voshall & Moriyama, 2018). To obtain the coverage of full-length proteins in an assembly, BLASTX v2.9.0 (Altschul et al., 1990) searches (e-value < 1e-20) of the assembled transcripts were performed against all known proteins in the Swissprot/Uniprot database (The UniProt Consortium et al., 2021). A full-length protein was reported in an assembly if a BLASTX match covered at least 90% of the protein’s length.

Another reference-dependent evaluation method is scanning the assembly for benchmarking universal single copy orthologs (BUSCOs) (Waterhouse et al., 2018). BUSCOs are expected to be highly conserved, therefore representing a known set of genes even in unknown organismic groups (Simão et al., 2015; Waterhouse et al., 2018). BUSCO recovery rates were obtained using the BUSCO v4 software and database (Manni et al., 2021). For *G. fossarum*, the arthropod BUSCO reference data set was used, comprising 1,013 BUSCOs. The insect reference BUSCO data set used for *E. danica* contained 1,367 BUSCOs.

To explore the consistency of expression estimates obtained from RSEM and Salmon, the Pearson correlation coefficient between their generated count data sets was calculated. Only homopolymer trimmed Trinity and rnaSPAdes data sets were included here. Since the count data sets include for each sample all transcripts, a correlation of the expression estimates of all transcripts was not possible due to the large number of comparisons (i.e. no. of genes x no. of samples). Therefore, we randomly selected 100 transcripts for this correlation analysis.

### Comparing the De Novo Pipelines With a Genome-Guided Pipeline

All selected evaluation metrics allow conclusions to be drawn about the quality of the generated transcriptomes and, consequently, the effectiveness of the chosen algorithm. Yet, the true transcriptome and transcript abundances remain unknown, and assembly and quantification tools rely on assumptions (see Supplementary Material Section S2, S3). In general, genome-guided transcript reconstruction and quantification can be more accurate because a reference genome mitigates the challenges associated with the assembly of highly uneven coverage of expression data: the genome-guided transcript reconstruction allows to resolve ambiguities, resulting in less redundant transcriptomes, and more reads being uniquely assigned to their genomic origin during abundance estimation. Since a reference genome is available for *E. danica* (NCBI BioProject PRJNA171755), we compared expression estimates obtained from the *de novo* approach with those obtained from a genome-guided analysis. The aim of this comparison was to assess the ability of a *de novo* analysis pipeline to retrieve the expression signal reliably. Only count data sets that were homopolymer trimmed and generated with Trinity or rnaSPAdes were included in this comparison.

For the genome-guided analysis, homopolymer trimmed RNA-seq reads were aligned against the *E. danica* reference genome with HISAT2 v2.2.1 (Kim et al., 2019) as follows: first, splice sites and exons were extracted from the genomic feature annotation and the genome was indexed. Next, all libraries were individually mapped against the reference genome in strand-specific mode (--rna-strandness RF) with a maximum intron length of 350,000. Alignments were reported for downstream transcriptome assembly (--dta) and transformed to coordinate sorted bam files with samtools v1.10 (Danecek et al., 2021). Transcript reconstruction and expression estimation were conducted with Stringtie v2.2.1 (Pertea et al., 2015) in stranded library mode (--fr) with a minimum isoform fraction of 0.01. Final counts were derived from coverage estimates using the prepDE.py script, which is provided with Stringtie, using an average read length of 150 bp.

To infer the expression signal in the different count data sets (i.e., generated with the *de novo* pipelines and in genome-guided mode) in an unsupervised manner, the count data was normalized with DESeq2 v1.34.0 (Love et al., 2014) and principal component analyses (PCAs) were performed using the R package PCAtools v2.6.0 (Blighe & Lun, 2021). All bioinformatic processing steps were performed on a Linux based HPC server, in a Snakemake v7.20.0 (Köster & Rahmann, 2012) workflow. Visualization of results was conducted in RStudio v2022.07.2 (RStudio Team, 2022) with R v4.2.2 (R Core Team, 2022) using the R package ggplot2 v3.4.1 (Wickham, 2017) and ggridges v0.5.4 (Wilke, 2022).

## Results

When homopolymers were removed, slightly fewer bases and contigs were assembled (Tables 2, 3), presumably due to the marginally reduced numbers of input bases in these data sets.

**Table 2:** Basic statistics of the different assemblies generated with the *Gammarus fossarum* RNA-seq data. QT = Quality trimming, H + QT = Homopolymer removal + quality trimming, Mbp = mega base pairs.

| Assembler | Trimming strategy | Assembled bases [Mbp] | No. of contigs | Contig length [bp] |  |  |  | No. of full-length proteins |
| --- | --- | --- | --- | --- | --- | --- | --- | --- |
|  |  |  |  | Min. | Max. | Mean | Median |  |
| Trinity | QT | 998.5 | 1,393,435 | 180 | 33,568 | 717 | 376 | 9,171 |
|  | H + QT | 980.2 | 1,372,461 | 168 | 32,242 | 714 | 376 | 9,153 |
| rnaSPAdes | QT | 838.9 | 1,045,534 | 73 | 28,548 | 802 | 378 | 8,648 |
|  | H + QT | 817.1 | 1,025,640 | 73 | 28,545 | 797 | 380 | 8,689 |
| IDBA-tran | QT | 539.9 | 1,054,917 | 200 | 21,562 | 512 | 352 | 4,533 |
|  | H + QT | 532.9 | 1,043,527 | 200 | 21,562 | 511 | 352 | 4,504 |

**Table 3:** Basic statistics of the different assemblies generated with the *Ephemera danica* RNA-seq data. QT = Quality trimming, H + QT = Homopolymer removal + quality trimming, Mbp = mega base pairs.

| Assembler | Trimming strategy | Assembled bases [Mbp] | No. of contigs | Contig length [bp] |  |  |  | No. of full-length proteins |
| --- | --- | --- | --- | --- | --- | --- | --- | --- |
|  |  |  |  | Min. | Max. | Mean | Median |  |
| Trinity | QT | 743.7 | 1,142,363 | 174 | 32,439 | 651 | 367 | 8,628 |
|  | H + QT | 738.4 | 1,131,805 | 178 | 44,525 | 652 | 367 | 8,620 |
| rnaSPAdes | QT | 646.5 | 664,539 | 93 | 32,452 | 973 | 427 | 7,998 |
|  | H + QT | 634.9 | 649,827 | 186 | 32,452 | 977 | 429 | 7,963 |
| IDBA-tran | QT | 308.3 | 647,645 | 200 | 18,511 | 476 | 342 | 4,063 |
|  | H + QT | 305 | 636,905 | 200 | 18,511 | 479 | 344 | 4,066 |

While the shape of the ExN50 curve of Trinity assemblies slightly changed when the homopolymer removal step was performed (Fig. S1), the evaluation metrics were generally unaffected by the trimming strategy. However, the choice of the assembly software had a strong impact on the resulting transcriptomes: Trinity and rnaSPAdes generated assemblies with decent contiguity, reflected in ExN50 values peaking on average at 1,910/2,630 bp (Trinity/rnaSPAdes) for *G. fossarum* and at 2,056/2,756 bp (Trinity/rnaSPAdes) for *E. danica* assemblies. Even though rnaSPAdes produced the more contiguous assemblies, Trinity consistently reconstructed the largest number of contigs as well as transcripts corresponding to full-length proteins (Tables 2, 3).

Both, Trinity and rnaSPAdes assembled transcriptomes in which more than 96% complete BUSCOs were always recovered (Fig. 1). In contrast, IDBA-tran assemblies contained the fewest transcripts corresponding to full-length proteins (Tables 2, 3) but the largest numbers of missing and fragmented BUSCOs (Fig. 1), which is consistent with the overall lower contiguity of these assemblies, indicated by ExN50 values peaking on average at 864 bp and 712 bp for *G. fossarum* and *E. danica* assemblies, respectively. If the assembly quality was sufficiently high, i.e., assemblies were generated with Trinity and rnaSPAdes, RSEM and Salmon were able to identify the origin of at least 86% of the read data, as opposed to the low read remapping rates observed for IDBA-tran assemblies (consistently < 60%) (Fig. 2).

**Fig. 1:**
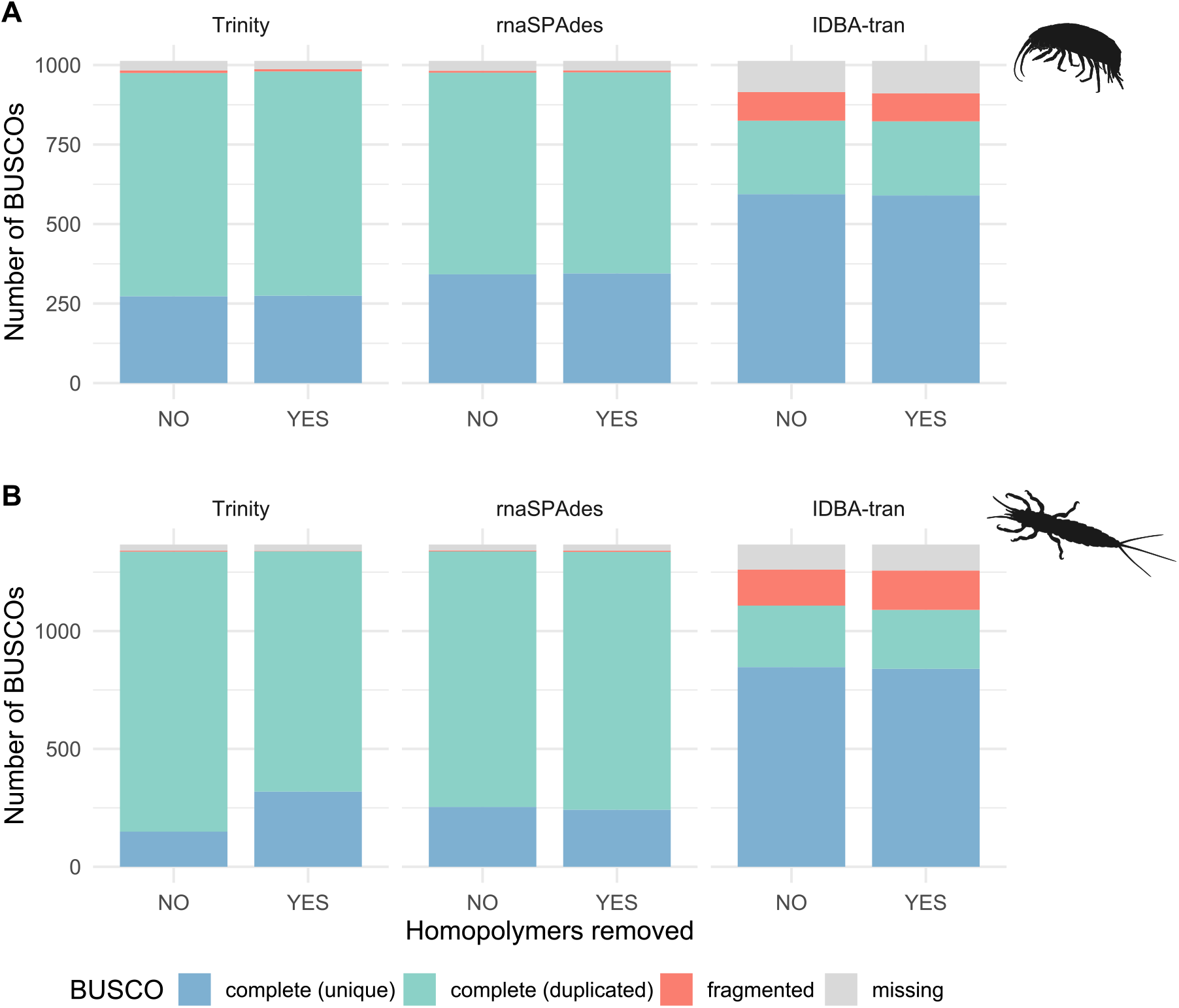
BUSCO recovery in *G. fossarum* (top) and *E. danica* (bottom) assemblies. Due to alternative splicing, transcriptome assemblies typically contain a large proportion of duplicated BUSCOs.

**Fig. 2:**
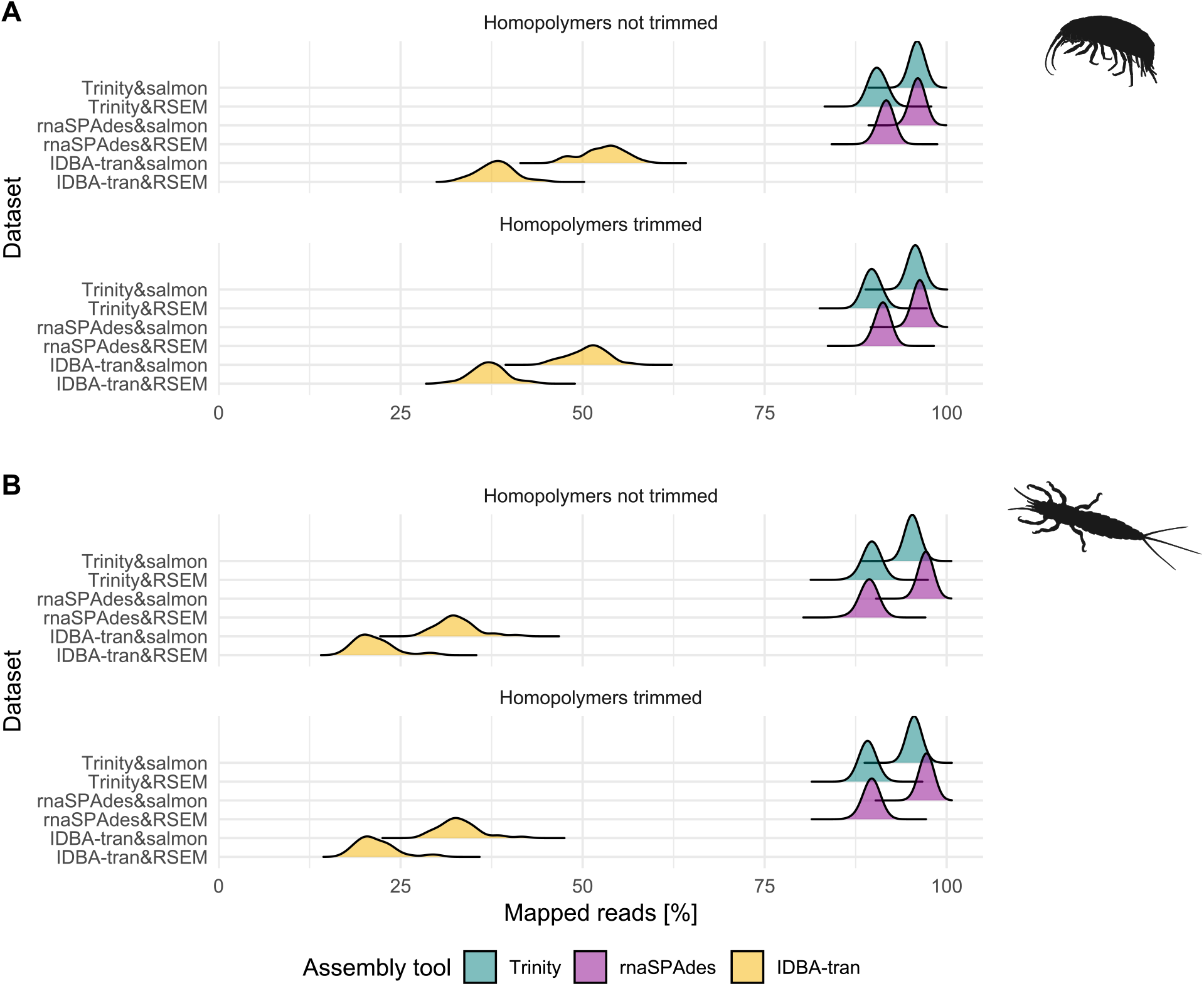
Density distributions of mapping rates of reads used to assemble the transcriptomes of *G. fossarum* (top) and *E. danica* (bottom).

Although Salmon produced slightly higher read mapping rates than RSEM (Fig. 2), the transcript counts estimated by quantification tools were overall congruent and highly correlated (Fig. 3); for the counts estimated for the *G. fossarum* data assembled with rnaSPAdes, RSEM estimated consistently higher abundances than Salmon (Fig. 3B).

**Fig. 3:**
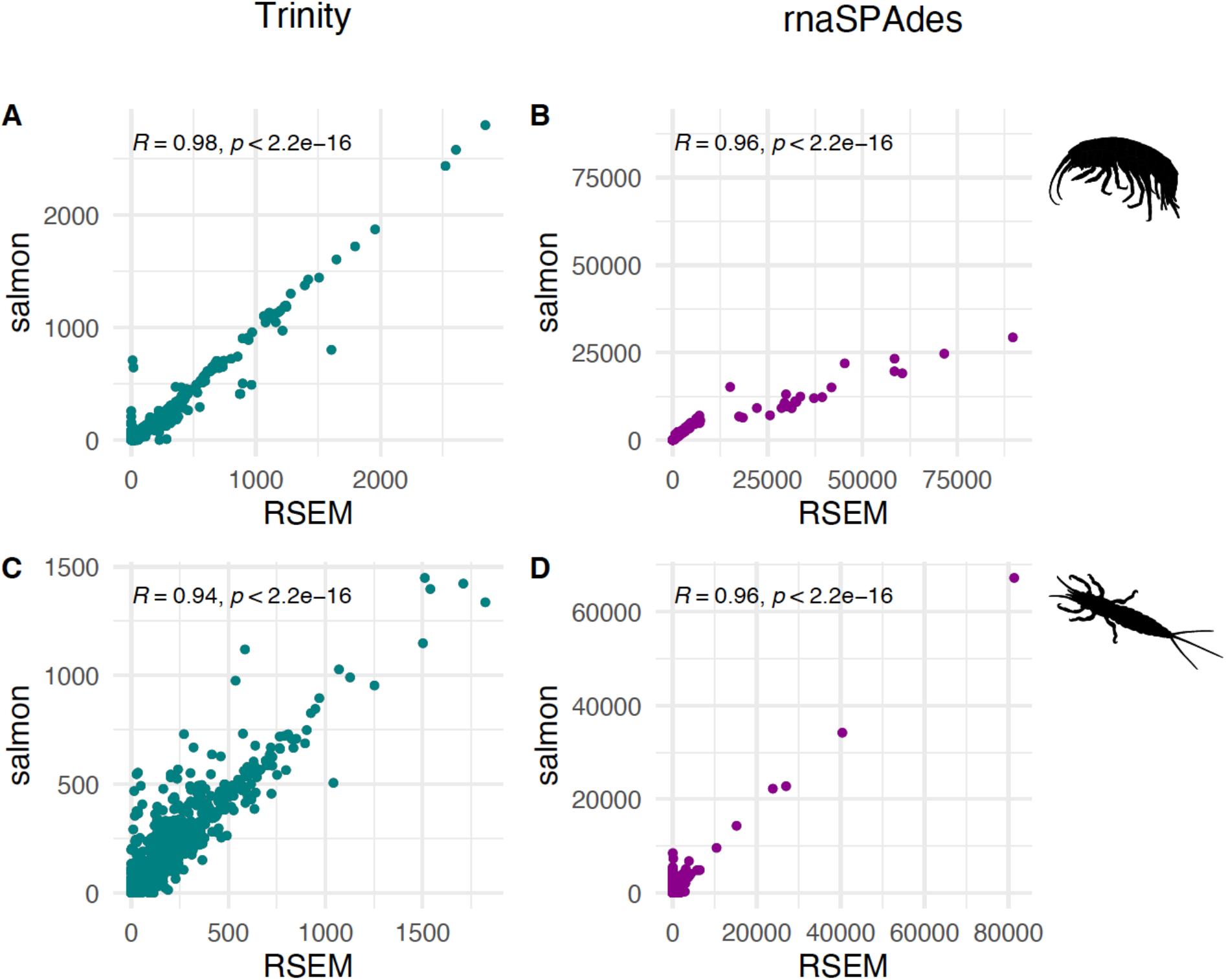
Correlation between counts estimated with RSEM and Salmon, for 100 randomly selected genes in 32 *G. fossarum* samples (A,B) and 36 *E. danica* samples (C,D). P-values were obtained from a t-test.

Finally, we found that the ordination-based inferences of the expression signal in the *E. danica* data were consistent between count data sets generated by the *de novo* pipelines and the count data generated using a genomic reference: although the orientation of the first two principal component axes can be inverted, all biplots depict the same relationship between the different RNA-seq libraries (Fig. 4).

**Fig. 4:**
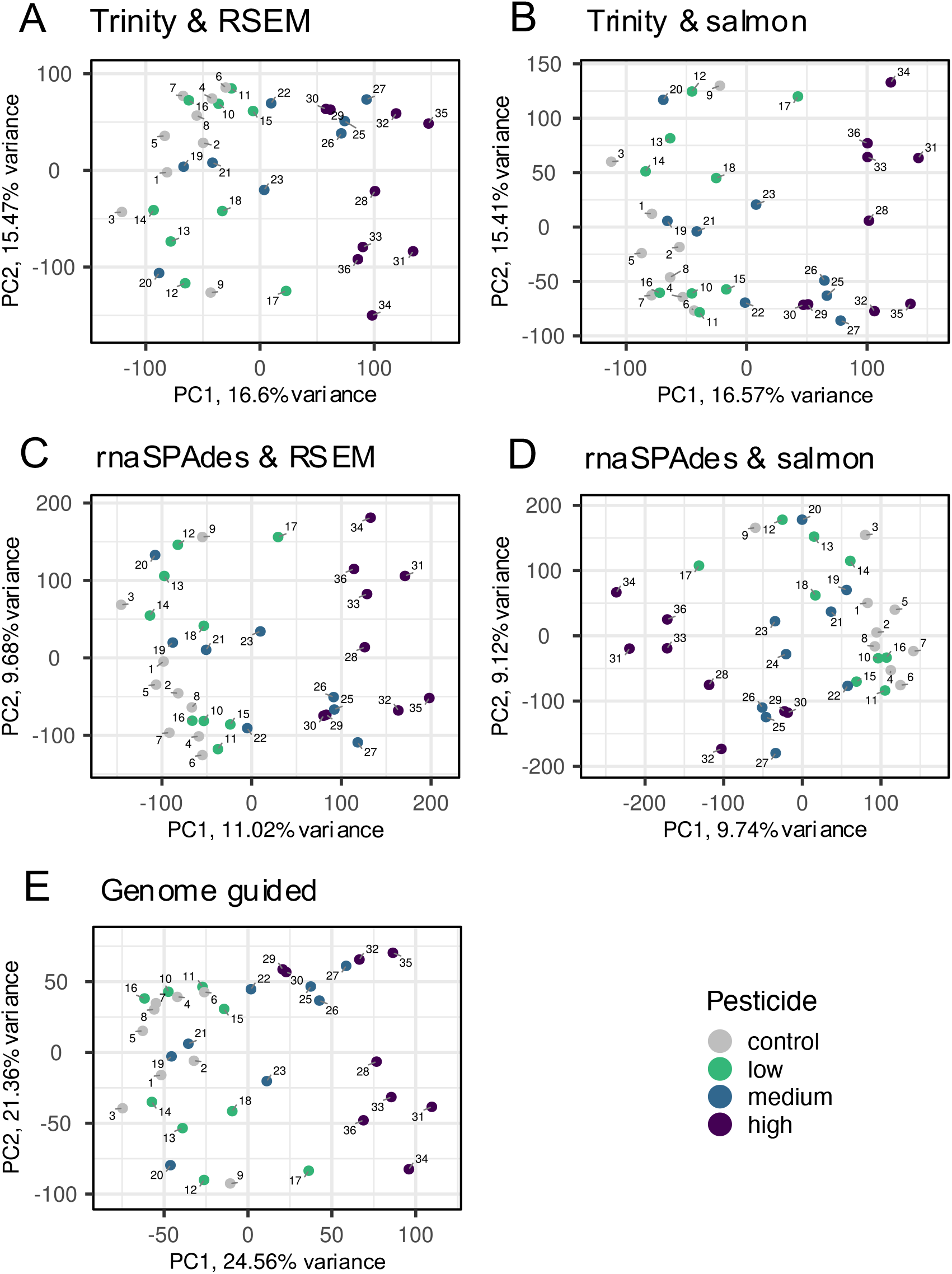
Principal component analyses based on the *E. danica* count data sets generated with *de novo* assembly pipelines (A-D) and the genome-guided transcript reconstruction and quantification (E). Due to lower redundancy in the expression data obtained from genome-guided bioinformatic processing, the first two principal components account for higher proportions of explained variance. This expression data set was obtained from an experiment in which *E. danica* was exposed to a pesticide in three different concentrations (low, medium, high). For further information see Brasseur et al., (2023).

## Discussion

Bioinformatic processing of non-model organism RNA-seq data for DGE analyses comprises three major steps: data pre-processing, *de novo* assembly and transcript quantification. We found that the strongest effect was, by far, introduced by the choice of the assembler (step 2). This observation is consistent across all data sets and metrics, whereas the applied trimming strategy and the choice of the quantification algorithms had no considerable impact.

### The Choice of the Trimming Approach

Theoretically, low-complexity reads can result in chimeric contigs, which can be misassembled based on artificially shared sequence information (Bushmanova et al., 2019), or introduce read mapping biases (Li et al., 2010). In practice, however, we could not identify any obvious benefits of the homopolymer removal based on the evaluation metrics. The observed variation in the ExN50 curves (Fig. S1) between the Trinity assemblies are likely rather related to the non-deterministic behavior of Trinity (Haas et al., 2013), than induced by the trimming approach. The impact of the trimming strategy is presumably limited because modern Illumina data are typically of high quality i.e., the majority of base quality (Phred) scores are above 30 (Illumina website, accessed on 29.09.2024), and because assembly programs correct for or discard low-complexity sequences (Bushmanova et al., 2019; Grabherr et al., 2011). However, a stronger impact of quality trimming might arise if the quality of the data is low or the sequencing depth is shallow. Therefore, we argue that it can be reasonable to remove technical artifacts such as homopolymers from the sequencing data to retain only biologically valid reads.

### The Choice of the Assembler

Based on the evaluation metrics, IDBA-tran appears to be unsuited for the assembly of RNA-seq data analyzed in this study. Here, we observed the lowest ExN50 values and the fewest full-length proteins as well as the highest number of incomplete and missing BUSCOs, indicating that these transcriptomes were more fragmented and less complete than assemblies generated with Trinity and rnaSPAdes. In fact, the low remapping rates suggest that IDBA-tran did not effectively leverage the input read data. The two other assemblers performed generally well on both species data sets, yet small differences regarding their sensitivity and specificity were observed that need to be considered.

The Trinity assemblies contained the highest number of transcripts corresponding to full-length proteins and complete BUSCOs, indicating that Trinity is more sensitive than rnaSPAdes. This sensitivity, however, comes at the expense of contiguity since rare (i.e, lowly expressed) transcripts are more likely to be partially assembled, decreasing the ExN50 value of Trinity assemblies. It should be noted that the principle of contiguity does not perfectly apply to transcriptome assemblies which are, per definition, dynamic and that larger (Ex)N50 values do not necessarily reflect the true underlying biology but potentially reward assemblers that over-assemble sequences (Voshall & Moriyama, 2018). However, its high sensitivity makes Trinity assemblies prone for inflation, i.e., it assembles by far the largest sets of contigs with often a substantial redundancy. For instance, Trinity reported nearly identical sequences as different isoforms, which only differed in length or by single nucleotide polymorphism. These might be technical artifacts, introduced by methodological imperfections such as incomplete stranding (Zeng & Mortazavi, 2012) or sequencing errors (Stoler & Nekrutenko, 2021), or represent biological redundancy. Freedman et al. (2021) showed that Trinity assembles allelic variants within and between organisms and reports these as different isoforms, a behavior which is especially important to consider if pooled RNA-seq data from wild-type (i.e., genetically diverse) organisms should be assembled. In these cases, an assembler such as rnaSPAdes, which performed best in terms of contiguity and produced less redundant assemblies, can be a more appropriate choice, although the lower sensitivity might come at the expense of missing rare transcripts.

Trinity and rnaSPAdes approach the trade-off between sensitivity and specificity by implementing different k-mer selection approaches. Trinity assembles the read data based on 25-mers (Grabherr et al., 2011). These short k-mers are beneficial to resolve rare transcripts because the required overlap is small. However, short k-mers are also more likely shared by unrelated contigs, thereby increasing the number of chimeras (Wang & Gribskov, 2017; Zhao et al., 2011). A larger value of k improves the assembly in terms of contiguity and correctness but comes at the cost of missing lowly expressed transcripts (Wang & Gribskov, 2017; Zhao et al., 2011). In rnaSPAdes, a robust assembly, derived from the larger k-mer size (i.e., k = 73 if read length = 150 bp), is extended by rare transcripts that were only resolved with the smaller k-mer size (i.e., k = 49 if read length = 150 bp). As such, the multi k-mer approach of rnaSPAdes aids in retrieving the full transcriptomic spectrum (Chen et al., 2011; Zhao et al., 2011) but Trinity was still found to be more sensitive. The final choice of the assembler might further depend on other characteristics and, depending on the data set, post-assembly improvement might be necessary to e.g., reduce redundancy such as assembly thinning or clustering (for further advice, see Raghavan et al., 2022).

### The Choice of the Quantification Method

While remapping rates primarily indicate how well the assembler exploited the input read data, they also inform about the ability of the quantification tool to identify the origin of reads in the reference. The slightly higher read mapping rates of Salmon compared to RSEM indicate that Salmon is less strict when defining read mapping thresholds. High remapping rates are important since the aim is to leverage the read data as efficiently as possible. However, an accurate quantification should be preferred over an overly permissive one. We think that the strong agreement between both tools, indicated by the high correlation of estimated counts, is a good indicator for an accurate quantification, although the tools estimated different absolute transcript abundances. While it is not possible to assess which of the derived counts are more correct, a systematic over- or underestimation of the ‘true’ count is presumably less of concern for DGE analyses, which focus on relative expression changes (i.e., fold change between samples). Important to note is that, although we did not observe any qualitative differences in the performance of both quantification programs, Salmon requires significantly less computing time (e.g., for the *E. danica* data set ∼ 2 days) than RSEM (e.g., for the *E. danica* data set ∼ 10 days). This highly increased efficiency of Salmon is not only enabled by its alignment-free mapping approach, but also by a reduction of the data complexity: while the initial inference problem is represented in the read data space, Salmon groups reads based on alignment information in equivalence classes (Patro et al., 2017). As such, the number of classes grows with the complexity of the transcriptome, not with the number of input reads. Considering that RNA-seq data sets become increasingly complex, researchers will favor computational solutions which can scale with the amount of data produced.

## Conclusion

Establishing an appropriate bioinformatic processing pipeline for RNA-seq data from non-model organisms can be challenging because the performance of computational algorithms varies between different data sets. In particular, the choice of the assembler can have a strong impact on the quality of the resulting transcriptome assembly, whereas the quality trimming strategy and the choice of the quantification tool seems to be less influential. Since the performance of assemblers can be highly unsatisfactory, as in the case of IDBA-tran, several assemblies should be generated with different programs and evaluated. However, the true underlying transcriptome remains unknown and while evaluation metrics can be used to identify assemblies of poor quality, the distinction between ‘good’ assemblies is not straightforward. In this regard, it is reassuring that, despite the numerous approximations made by transcriptome assemblers and transcript quantification algorithms, the retrieved biological signal is consistent between count data sets generated by the different *de novo* pipelines and the count data generated using a genomic reference. This implies that, although we are currently far away from a scenario in which high-quality genomes are accessible for most taxonomic groups, the *de novo* approach is a valid and robust analysis strategy for RNA-seq data sets obtained from non-model organisms.

## References

Adams, M. D., Celniker, S. E., Holt, R. A., Evans, C. A., Gocayne, J. D., Amanatides, P. G.,Scherer, S. E., Li, P. W., Hoskins, R. A., Galle, R. F., George, R. A., Lewis, S. E., Richards, S., Ashburner, M., Henderson, S. N., Sutton, G. G., Wortman, J. R., Yandell, M. D., Zhang, Q., … Venter, J. C. (2000). The Genome Sequence of Drosophila melanogaster. Science, 287(5461), 2185–2195. 10.1126/science.287.5461.2185

Altschul, S. F., Gish, W., Miller, W., Myers, E. W., & Lipman, D. J. (1990). Basic local alignment search tool. Journal of Molecular Biology, 215(3), 403–410. 10.1016/S0022-2836(05)80360-2

Andrews, S. (2010). FastQC A Quality Control tool for High Throughput Sequence Data. https://www.bioinformatics.babraham.ac.uk/projects/fastqc/

Andrews, S. (2016, May 4). QC Fail Sequencing » Illumina 2 colour chemistry can overcall high confidence G bases. https://sequencing.qcfail.com/articles/illumina-2-colour-chemistry-can-overcall-high-confidence-g-bases/

Blighe, K., & Lun, A. (2021). PCAtools: Everything principal components analysis. https://github.com/kevinblighe/PCAtools

Brasseur, M. V., Beermann, A. J., Elbrecht, V., Grabner, D., Peinert-Voss, B., Salis, R., Weiss, M., Mayer, C., & Leese, F. (2022). Impacts of multiple anthropogenic stressors on the transcriptional response of Gammarus fossarum in a mesocosm field experiment. BMC Genomics, 23(1), 816. 10.1186/s12864-022-09050-1

Brasseur, M. V., Leese, F., Schäfer, R. B., Schreiner, V. C., & Mayer, C. (2023). Transcriptomic sequencing data illuminate insecticide-induced physiological stress mechanisms in aquatic non-target invertebrates. Environmental Pollution, 335, 122306. 10.1016/j.envpol.2023.122306

Bray, N. L., Pimentel, H., Melsted, P., & Pachter, L. (2016). Near-optimal probabilistic RNA-seq quantification. Nature Biotechnology, 34(5), Article 5. 10.1038/nbt.3519

Bushmanova, E., Antipov, D., Lapidus, A., & Prjibelski, A. D. (2019). rnaSPAdes: A de novo transcriptome assembler and its application to RNA-Seq data. GigaScience, 8(9), giz100. 10.1093/gigascience/giz100

Chandramohan, R., Po-Yen Wu, Phan, J. H., & Wang, M. D. (2013). Benchmarking RNA-Seq quantification tools. 2013 35th Annual International Conference of the IEEE Engineering in Medicine and Biology Society (EMBC), 647–650. 10.1109/EMBC.2013.6609583

Chen, G., Yin, K., Wang, C., & Shi, T. (2011). De novo transcriptome assembly of RNA-Seq reads with different strategies. Science China Life Sciences, 54(12), 1129–1133. 10.1007/s11427-011-4256-9

Clarke, K., Yang, Y., Marsh, R., Xie, L., & K., Z. K. (2013). Comparative analysis of de novo transcriptome assembly. Science China Life Sciences, 56(2), 156–162. 10.1007/s11427-013-4444-x

Conesa, A., Madrigal, P., Tarazona, S., Gomez-Cabrero, D., Cervera, A., McPherson, A., Szcześniak, M. W., Gaffney, D. J., Elo, L. L., Zhang, X., & Mortazavi, A. (2016). A survey of best practices for RNA-seq data analysis. Genome Biology, 17(1), 13. 10.1186/s13059-016-0881-8

Danecek, P., Bonfield, J. K., Liddle, J., Marshall, J., Ohan, V., Pollard, M. O., Whitwham, A., Keane, T., McCarthy, S. A., Davies, R. M., & Li, H. (2021). Twelve years of SAMtools and BCFtools. GigaScience, 10(2), giab008. 10.1093/gigascience/giab008

Ekblom, R., & Galindo, J. (2011). Applications of next generation sequencing in molecular ecology of non-model organisms. Heredity, 107(1), Article 1. 10.1038/hdy.2010.152

Eldem, V., Zararsiz, G., TaŞçi, T., Duru, I. P., Bakir, Y., & Erkan, M. (2017). Transcriptome Analysis for Non-Model Organism: Current Status and Best-Practices. In F. A. Marchi, P. D. R. Cirillo, & E. C. Mateo (Eds.), Applications of RNA-Seq and Omics Strategies— From Microorganisms to Human Health. InTech. 10.5772/intechopen.68983

Fabbro, C. D., Scalabrin, S., Morgante, M., & Giorgi, F. M. (2013). An Extensive Evaluation of Read Trimming Effects on Illumina NGS Data Analysis. PLOS ONE, 8(12), e85024. 10.1371/journal.pone.0085024

Finotello, F., & Di Camillo, B. (2015). Measuring differential gene expression with RNA-seq: Challenges and strategies for data analysis. Briefings in Functional Genomics, 14(2), 130–142. 10.1093/bfgp/elu035

Fox, E. J., Reid-Bayliss, K. S., Emond, M. J., & Loeb, L. A. (2014). Accuracy of Next Generation Sequencing Platforms. Next Generation, Sequencing & Applications, 1, 1000106. 10.4172/jngsa.1000106

Freedman, A. H., Clamp, M., & Sackton, T. B. (2021). Error, noise and bias in de novo transcriptome assemblies. Molecular Ecology Resources, 21(1), 18–29. 10.1111/1755-0998.13156

García-Nieto, P. E., Wang, B., & Fraser, H. B. (2022). Transcriptome diversity is a systematic source of variation in RNA-sequencing data. PLOS Computational Biology, 18(3), e1009939. 10.1371/journal.pcbi.1009939

Geniza, M., & Jaiswal, P. (2017). Tools for building de novo transcriptome assembly. Current Plant Biology, 11–12, 41–45. 10.1016/j.cpb.2017.12.004

Grabherr, M. G., Haas, B. J., Yassour, M., Levin, J. Z., Thompson, D. A., Amit, I., Adiconis, X., Fan, L., Raychowdhury, R., Zeng, Q., Chen, Z., Mauceli, E., Hacohen, N., Gnirke, A., Rhind, N., di Palma, F., Birren, B. W., Nusbaum, C., Lindblad-Toh, K., … Regev, A. (2011). Trinity: Reconstructing a full-length transcriptome without a genome from RNA-Seq data. Nature Biotechnology, 29(7), 644–652. 10.1038/nbt.1883

Haas, B. J., Papanicolaou, A., Yassour, M., Grabherr, M., Blood, P. D., Bowden, J., Couger, M. B., Eccles, D., Li, B., Lieber, M., MacManes, M. D., Ott, M., Orvis, J., Pochet, N., Strozzi, F., Weeks, N., Westerman, R., William, T., Dewey, C. N., … Regev, A. (2013). De novo transcript sequence reconstruction from RNA-seq using the Trinity platform for reference generation and analysis. Nature Protocols, 8(8), Article 8. 10.1038/nprot.2013.084

Hölzer, M., & Marz, M. (2019). De novo transcriptome assembly: A comprehensive cross-species comparison of short-read RNA-Seq assemblers. GigaScience, 8(5). 10.1093/gigascience/giz039

Kanitz, A., Gypas, F., Gruber, A. J., Gruber, A. R., Martin, G., & Zavolan, M. (2015). Comparative assessment of methods for the computational inference of transcript isoform abundance from RNA-seq data. Genome Biology, 16(1), 150. 10.1186/s13059-015-0702-5

Kim, D., Paggi, J. M., Park, C., Bennett, C., & Salzberg, S. L. (2019). Graph-based genome alignment and genotyping with HISAT2 and HISAT-genotype. Nature Biotechnology, 37(8), Article 8. 10.1038/s41587-019-0201-4

Köster, J., & Rahmann, S. (2012). Snakemake—A scalable bioinformatics workflow engine. Bioinformatics, 28(19), 2520–2522. 10.1093/bioinformatics/bts480

Langmead, B., & Salzberg, S. L. (2012). Fast gapped-read alignment with Bowtie 2. Nature Methods, 9(4), Article 4. 10.1038/nmeth.1923

Li, B., & Dewey, C. N. (2011). RSEM: Accurate transcript quantification from RNA-Seq data with or without a reference genome. BMC Bioinformatics, 12(1), 323. 10.1186/1471-2105-12-323

Li, B., Ruotti, V., Stewart, R. M., Thomson, J. A., & Dewey, C. N. (2010). RNA-Seq gene expression estimation with read mapping uncertainty. Bioinformatics, 26(4), 493–500. 10.1093/bioinformatics/btp692

Love, M. I., Huber, W., & Anders, S. (2014). Moderated estimation of fold change and dispersion for RNA-seq data with DESeq2. Genome Biology, 15(12), 550. 10.1186/s13059-014-0550-8

Lu, B., Zeng, Z., & Shi, T. (2013). Comparative study of de novo assembly and genome-guided assembly strategies for transcriptome reconstruction based on RNA-Seq. Science China Life Sciences, 56(2), 143–155. 10.1007/s11427-013-4442-z

MacManes, M. D. (2014). On the optimal trimming of high-throughput mRNA sequence data. Frontiers in Genetics, 5. 10.3389/fgene.2014.00013

Manni, M., Berkeley, M. R., Seppey, M., Simão, F. A., & Zdobnov, E. M. (2021). BUSCO update: Novel and streamlined workflows along with broader and deeper phylogenetic coverage for scoring of eukaryotic, prokaryotic, and viral genomes. Molecular Biology and Evolution, 38(10), 4647–4654. 10.1093/molbev/msab199

Martin, M. (2011). Cutadapt removes adapter sequences from high-throughput sequencing reads. EMBnet.Journal, 17(1), 10–12. 10.14806/ej.17.1.200

Mortazavi, A., Williams, B. A., McCue, K., Schaeffer, L., & Wold, B. (2008). Mapping and quantifying mammalian transcriptomes by RNA-Seq. Nature Methods, 5(7), 621–628. 10.1038/nmeth.1226

Nariai, N., Kojima, K., Mimori, T., Sato, Y., Kawai, Y., Yamaguchi-Kabata, Y., & Nagasaki, M. (2014). TIGAR2: Sensitive and accurate estimation of transcript isoform expression with longer RNA-Seq reads. BMC Genomics, 15(10), S5. 10.1186/1471-2164-15-S10-S5

Nicolae, M., Mangul, S., Măndoiu, I. I., & Zelikovsky, A. (2011). Estimation of alternative splicing isoform frequencies from RNA-Seq data. Algorithms for Molecular Biology, 6(1), 9. 10.1186/1748-7188-6-9

Patro, R., Duggal, G., Love, M. I., Irizarry, R. A., & Kingsford, C. (2017). Salmon provides fast and bias-aware quantification of transcript expression. Nature Methods, 14(4), Article 4. 10.1038/nmeth.4197

Patro, R., Mount, S. M., & Kingsford, C. (2014). Sailfish enables alignment-free isoform quantification from RNA-seq reads using lightweight algorithms. Nature Biotechnology, 32(5), Article 5. 10.1038/nbt.2862

Peng, Y., Leung, H. C. M., Yiu, S.-M., Lv, M.-J., Zhu, X.-G., & Chin, F. Y. L. (2013). IDBA-tran: A more robust de novo de Bruijn graph assembler for transcriptomes with uneven expression levels. Bioinformatics, 29(13), i326–i334. 10.1093/bioinformatics/btt219

Pertea, M., Pertea, G. M., Antonescu, C. M., Chang, T.-C., Mendell, J. T., & Salzberg, S. L. (2015). StringTie enables improved reconstruction of a transcriptome from RNA-seq reads. Nature Biotechnology, 33(3), Article 3. 10.1038/nbt.3122

Pickrell, J. K., Marioni, J. C., Pai, A. A., Degner, J. F., Engelhardt, B. E., Nkadori, E., Veyrieras, J.-B., Stephens, M., Gilad, Y., & Pritchard, J. K. (2010). Understanding mechanisms underlying human gene expression variation with RNA sequencing. Nature, 464(7289), Article 7289. 10.1038/nature08872

R Core Team. (2022). R: a language and environment for statistical computing. R Foundation for Statistical Computing. https://www.R-project.org/

Raghavan, V., Kraft, L., Mesny, F., & Rigerte, L. (2022). A simple guide to de novo transcriptome assembly and annotation. Briefings in Bioinformatics, 23(2), bbab563. 10.1093/bib/bbab563

RStudio Team. (2022). RStudio: Integrated Development Environment for R. RStudio, PBC. http://www.rstudio.com/

Schröder, J., Bailey, J., Conway, T., & Zobel, J. (2010). Reference-Free Validation of Short Read Data. PLoS ONE, 5(9), e12681. 10.1371/journal.pone.0012681

Shi, H., Zhou, Y., Jia, E., Pan, M., Bai, Y., & Ge, Q. (2021). Bias in RNA-seq Library Preparation: Current Challenges and Solutions. BioMed Research International, 2021, e6647597. 10.1155/2021/6647597

Simão, F. A., Waterhouse, R. M., Ioannidis, P., Kriventseva, E. V., & Zdobnov, E. M. (2015). BUSCO: Assessing genome assembly and annotation completeness with single-copy orthologs. Bioinformatics, 31(19), 3210–3212. 10.1093/bioinformatics/btv351

Srivastava, A., Sarkar, H., Gupta, N., & Patro, R. (2016). RapMap: A rapid, sensitive and accurate tool for mapping RNA-seq reads to transcriptomes. Bioinformatics, 32(12), i192–i200. 10.1093/bioinformatics/btw277

Stoler, N., & Nekrutenko, A. (2021). Sequencing error profiles of Illumina sequencing instruments. NAR Genomics and Bioinformatics, 3(1). 10.1093/nargab/lqab019

Teng, M., Love, M. I., Davis, C. A., Djebali, S., Dobin, A., Graveley, B. R., Li, S., Mason, C. E., Olson, S., Pervouchine, D., Sloan, C. A., Wei, X., Zhan, L., & Irizarry, R. A. (2016). A benchmark for RNA-seq quantification pipelines. Genome Biology, 17(1), 74. 10.1186/s13059-016-0940-1

The UniProt Consortium, Bateman, A., Martin, M.-J., Orchard, S., Magrane, M., Agivetova, R., Ahmad, S., Alpi, E., Bowler-Barnett, E. H., Britto, R., Bursteinas, B., Bye-A-Jee, H., Coetzee, R., Cukura, A., Da Silva, A., Denny, P., Dogan, T., Ebenezer, T., Fan, J., … Teodoro, D. (2021). UniProt: The universal protein knowledgebase in 2021. Nucleic Acids Research, 49(D1), D480–D489. 10.1093/nar/gkaa1100

Van Pelt-Verkuil, E., Van Leeuwen, W. B., & Te Witt, R. (Eds.). (2019). Molecular Diagnostics: Part 1: Technical Backgrounds and Quality Aspects. Springer Singapore. 10.1007/978-981-13-1604-3

Voshall, A., & Moriyama, E. N. (2018). Next-Generation Transcriptome Assembly: Strategies and Performance Analaysis. In I. Y. Abdurakhmonov (Ed.), Bioinformatics in the Era of Post Genomics and Big Data. InTech. 10.5772/intechopen.73497

Wang, S., & Gribskov, M. (2017). Comprehensive evaluation of de novo transcriptome assembly programs and their effects on differential gene expression analysis. Bioinformatics, 33(3), 327–333. 10.1093/bioinformatics/btw625

Wang, Z., Gerstein, M., & Snyder, M. (2009). RNA-Seq: A revolutionary tool for transcriptomics. Nature Reviews Genetics, 10(1), Article 1. 10.1038/nrg2484

Waterhouse, R. M., Seppey, M., Simão, F. A., Manni, M., Ioannidis, P., Klioutchnikov, G., Kriventseva, E. V., & Zdobnov, E. M. (2018). BUSCO Applications from Quality Assessments to Gene Prediction and Phylogenomics. Molecular Biology and Evolution, 35(3), 543–548. 10.1093/molbev/msx319

Wickham, H. (2017). ggplot2: Elegant graphics for data analysis (2nd Edition). Journal of Statistical Software, 77(Book Review 2). 10.18637/jss.v077.b02

Wilke, C. O. (2022). ggridges: Ridgeline Plots in “ggplot2.” https://wilkelab.org/ggridges/

Williams, C. R., Baccarella, A., Parrish, J. Z., & Kim, C. C. (2016). Trimming of sequence reads alters RNA-Seq gene expression estimates. BMC Bioinformatics, 17(1), 103. 10.1186/s12859-016-0956-2

Zeng, W., & Mortazavi, A. (2012). Technical considerations for functional sequencing assays. Nature Immunology, 13(9), Article 9. 10.1038/ni.2407

Zhang, C., Zhang, B., Lin, L.-L., & Zhao, S. (2017). Evaluation and comparison of computational tools for RNA-seq isoform quantification. BMC Genomics, 18(1), 583. 10.1186/s12864-017-4002-1

Zhang, C., Zhang, B., Vincent, M. S., & Zhao, S. (2016). Bioinformatics Tools for RNA-seq Gene and Isoform Quantification. Journal of Next Generation Sequencing & Applications, 03(03). 10.4172/2469-9853.1000140

Zhao, Q.-Y., Wang, Y., Kong, Y.-M., Luo, D., Li, X., & Hao, P. (2011). Optimizing de novo transcriptome assembly from short-read RNA-Seq data: A comparative study. BMC Bioinformatics, 12(14), S2. 10.1186/1471-2105-12-S14-S2

