## Supplementary Material for "Comparative Analysis of De Novo Assemblers and Quantification Software for RNA-sequencing Data in Non-Model Arthropods"

**Supplementary Material S1 – homopolymer trimming**

Homopolymers can arise due to several reasons in RNA-Seq data sets: they might be a result of the two-color chemistry used by Illumina NovaSeq and NextSeq platforms, which interpret the absence of a fluorescence emission signal as the presence of the nucleotide guanine (G) (Van Pelt-Verkuil et al., 2019). These bases, which are dominantly inserted at the 3' end typically get very high confidence scores and may obscure the quality profile used by trimming tools, hindering the effective removal of lower-quality bases (Andrews, 2016). Further, poly-T or poly-A stretches can be found in RNA-Seq libraries enriched for mRNA by oligo(dT) priming. Several trimming tools can account for such technical artifacts: for instance, AfterQC (Chen et al., 2017) filters reads with abnormal PolyA/PolyT/PolyC/PolyG sequences, fastp (Chen et al., 2018) trims homopolymer tails at the reads 3' end and cutadapt (M. Martin, 2011) allows to remove poly-A tails from the forward read and poly-T "heads" from the reverse read. However, AfterQC only removes entire reads, whereas is successor fastp only trims perfect mono repeats of a specified length from the 3' end. In other words, fastp will not trim homopolymers beyond the specified length or homopolymers in the middle of a read. Cutadapt is more flexible in this regard, but limited to trim A-/T-stretches, at least in its homopolymer removal mode. Therefore, we developed a C++ program, which can be used to trim (unperfect) mono repeat stretches of all bases at the 3' end as well as the middle of the read prior to quality trimming.

**Supplementary material S2 – transcriptome assemblers**

Trinity (Grabherr et al., 2011), rnaSPAdes (Bushmanova et al., 2019) and IDBA-tran (Peng et al., 2013) use the *de Bruijn* graph assembly approach for sequencing reads, which can be essentially divided in the following steps (J. A. Martin & Wang, 2011):

1. Decomposing the reads into seeds with a fixed length k (k-mer). Unique k-mers represent the *de Bruijn* graph nodes.

2. Consecutively overlapping k-mers are connected by edges (overlap = k-1).

3. Variable sites induce branching of the graph structure.

4. Contigs are derived by traversing all possible path combinations through the *de Bruijn* graph.

5. Contigs that are well supported by the read data are reported as assembled isoforms.

Although all assemblers implement the same fundamental strategy, each program relies on different assumptions and thresholds to construct, correct, and traverse the *de Bruijn* graph.

Trinity involves three functional submodules, Inchworm, Chrysalis and Butterfly (see Grabherr et al., 2011). Inchworm creates a k-mer catalog (with k = 25), followed by the reconstruction of linear contigs based on overlapping (k-1)-mers through a greedy extension. The Inchworm module reports only the full-length sequence for a dominant isoform, but related contigs share sequence information by partially overlapping k-mers. Chrysalis clusters related contigs and constructs a *de Brujin* graph for each cluster. Butterfly operates on these graphs, collapsing the unbranched parts and identifying the path through the graph which is best supported by the read data. The construction of thousands of *de Bruijn* graphs, each ideally representing one gene and its transcriptional complexity, is a specific trait of Trinity, which was directly manufactured for RNA-sequencing data.

In contrast, rnaSPAdes and IDBA-tran, which are extensions of the genome assemblers SPAdes (Bankevich et al., 2012) and IDBA (Peng et al., 2010), respectively, build a single *de Bruijn* graph that is subsequently decomposed (Bushmanova et al., 2019; Peng et al., 2013). In rnaSPAdes, the construction of the assembly graph is followed by a simplification step, including repeat resolution and scaffolding based on read alignments to the graph (see Bushmanova et al., 2019). rnaSPAdes relies on an iterative assembly graph construction based on two values of k, which are dynamically determined based on the read length of the input data. The multi k-mer strategy is similarly adopted by IDBA-tran, which starts with a small k to construct a graph that is iteratively updated by graphs formed based on larger k (see Peng et al., 2013). In contrast to rnaSPAdes, the default values of k are fixed: the version of IDBA-tran used in this study initializes the assembly with $k_{min}$ = 20, increases k by increments of 10 in each iteration, until $k_{max}$ = 60. During each iteration, the previously generated contigs are treated as input reads for the following graph construction. Based on coverage information, the *de Bruijn* graph is modularized in components containing related contigs. A feature of IDBA-tran is the implementation of a probabilistic framework to model the error probability of a k-mer or a contig. Through progressive removal of erroneous contigs, connected subcomponents i.e., isoforms from the same gene, are finally resolved.

**Supplementary Material S3 – transcript quantification software**

Salmon (Patro et al., 2017) and RSEM (Li & Dewey, 2011) both frame the expression estimation problem in a maximum likelihood context. Apart from the expression parameter, their models implement parameters to correct for the non-uniform distribution of reads along a transcript, which arise during the generation of RNA-sequencing data e.g., due to differences in sequence composition and fragment length distribution. Further, both rely on the expectation-maximization (EM) algorithm (Dempster et al., 1977) to iteratively assign reads to transcripts, proportional to the current transcript abundances (Li & Dewey, 2011; Patro et al.,

2017). RSEM estimates the probability that a read derived from a specific transcript given the observed read alignment data (see Li & Dewey, 2011). In other words, RSEM estimates transcript abundances which maximize the likelihood of the estimated expression values (Li et al., 2010; Li & Dewey, 2011). During iterations of the EM algorithm, fractions of reads are proportionally assigned to their possible origins according to the relative expression of a transcript (expectation step). Then, the relative expression of a transcript is re-estimated based on the allocated read count (maximization step).

In the likelihood model of salmon, the expression parameter is defined as nucleotide fractions, which depend on transcript abundances (Li et al., 2010; Patro et al., 2017). Salmon aims to identify the nucleotide fractions of transcripts which maximize the probability that the nucleotides were sampled from a particular transcript given the expression data (see Patro et al., 2017). The bias model of salmon, which accounts for the non-uniform distribution of reads along a transcript, computes the probability that a particular read was generated given the specific transcript sequence i.e., its composition. During iterations of the EM-algorithm, read counts which are weighted by the bias model are proportionally assigned to the transcripts from which they were putatively sampled until model convergence.

### Supplementary material S4

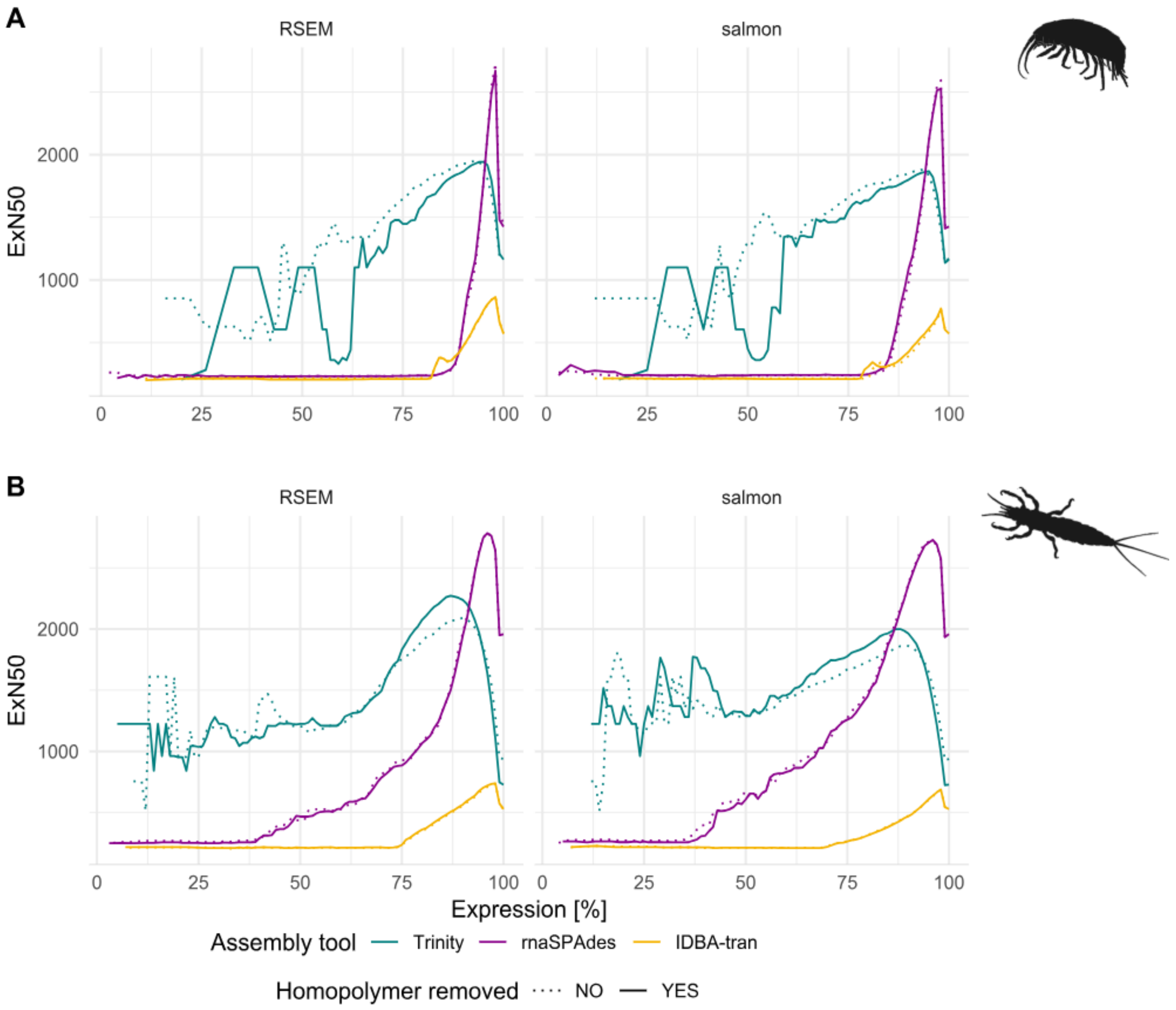


Figure S1: ExN50 values of *G. fossarum* (top) and *E. danica* (bottom) assemblies, obtained from abundance estimation using RSEM (left) and salmon (right). These transcripts are expected to be well supported by the read data and therefore accurately assembled. The different trimming strategies are indicated by dashed and solid lines. The different colors indicate the assembly program.

**Bibliography**


Andrews, S. (2016, May 4). *QC Fail Sequencing » Illumina 2 colour chemistry can overcall high confidence G bases*. https://sequencing.qcfail.com/articles/illumina-2-colour-chemistry-can-overcall-high-confidence-g-bases/

Bankevich, A., Nurk, S., Antipov, D., Gurevich, A. A., Dvorkin, M., Kulikov, A. S., Lesin, V. M., Nikolenko, S. I., Pham, S., Prjibelski, A. D., Pyshkin, A. V., Sirotkin, A. V., Vyahhi, N., Tesler, G., Alekseyev, M. A., & Pevzner, P. A. (2012). SPAdes: A New Genome Assembly Algorithm and Its Applications to Single-Cell Sequencing. *Journal of Computational Biology*, *19*(5), 455–477. https://doi.org/10.1089/cmb.2012.0021

Bushmanova, E., Antipov, D., Lapidus, A., & Prjibelski, A. D. (2019). rnaSPAdes: A de novo transcriptome assembler and its application to RNA-Seq data. *GigaScience*, *8*(9), giz100. https://doi.org/10.1093/gigascience/giz100

Chen, S., Huang, T., Zhou, Y., Han, Y., Xu, M., & Gu, J. (2017). AfterQC: Automatic filtering, trimming, error removing and quality control for fastq data. *BMC Bioinformatics*, *18*(3), 80. https://doi.org/10.1186/s12859-017-1469-3

Chen, S., Zhou, Y., Chen, Y., & Gu, J. (2018). fastp: An ultra-fast all-in-one FASTQ preprocessor. *Bioinformatics*, *34*(17), i884–i890. https://doi.org/10.1093/bioinformatics/bty560

Dempster, A. P., Laird, N. M., & Rubin, D. B. (1977). Maximum Likelihood from Incomplete Data via the EM Algorithm. *Journal of the Royal Statistical Society. Series B (Methodological)*, *39*(1), 1–38.

Grabherr, M. G., Haas, B. J., Yassour, M., Levin, J. Z., Thompson, D. A., Amit, I., Adiconis, X., Fan, L., Raychowdhury, R., Zeng, Q., Chen, Z., Mauceli, E., Hacohen, N., Gnirke, A., Rhind, N., di Palma, F., Birren, B. W., Nusbaum, C., Lindblad-Toh, K., … Regev, A. (2011). Trinity: Reconstructing a full-length transcriptome without a genome from RNA-Seq data. *Nature Biotechnology*, *29*(7), 644–652. https://doi.org/10.1038/nbt.1883

Li, B., & Dewey, C. N. (2011). RSEM: Accurate transcript quantification from RNA-Seq data with or without a reference genome. *BMC Bioinformatics*, *12*(1), 323. https://doi.org/10.1186/1471-2105-12-323

Li, B., Ruotti, V., Stewart, R. M., Thomson, J. A., & Dewey, C. N. (2010). RNA-Seq gene expression estimation with read mapping uncertainty. *Bioinformatics*, *26*(4), 493–500. https://doi.org/10.1093/bioinformatics/btp692

Martin, J. A., & Wang, Z. (2011). Next-generation transcriptome assembly. *Nature Reviews Genetics*, *12*(10), 671–682. https://doi.org/10.1038/nrg3068

Martin, M. (2011). Cutadapt removes adapter sequences from high-throughput sequencing reads. *EMBnet.Journal*, *17*(1), 10–12. https://doi.org/10.14806/ej.17.1.200

Patro, R., Duggal, G., Love, M. I., Irizarry, R. A., & Kingsford, C. (2017). Salmon provides fast and bias-aware quantification of transcript expression. *Nature Methods*, *14*(4), Article 4. https://doi.org/10.1038/nmeth.4197

Peng, Y., Leung, H. C. M., Yiu, S. M., & Chin, F. Y. L. (2010). IDBA – A Practical Iterative de Bruijn Graph De Novo Assembler. In B. Berger (Ed.), *Research in Computational Molecular Biology* (pp. 426–440). Springer. https://doi.org/10.1007/978-3-642-12683-3_28

Peng, Y., Leung, H. C. M., Yiu, S.-M., Lv, M.-J., Zhu, X.-G., & Chin, F. Y. L. (2013). IDBA-tran: A more robust de novo de Bruijn graph assembler for transcriptomes with uneven expression levels. *Bioinformatics*, *29*(13), i326–i334. https://doi.org/10.1093/bioinformatics/btt219

Van Pelt-Verkuil, E., Van Leeuwen, W. B., & Te Witt, R. (Eds.). (2019). *Molecular Diagnostics: Part 1: Technical Backgrounds and Quality Aspects*. Springer Singapore. https://doi.org/10.1007/978-981-13-1604-3
